# Microbowls with controlled concavity for accurate microscale mass spectrometry

**DOI:** 10.1101/2021.12.02.470972

**Authors:** Linfeng Xu, Xiangpeng Li, Wenzong Li, Kai-chun Chang, Hyunjun Yang, Nannan Tao, Pengfei Zhang, Emory Payne, Cyrus Modavi, Jacqueline Humphries, Chia-Wei Lu, Adam R. Abate

**Affiliations:** Department of Bioengineering and Therapeutic sciences, University of California, San Francisco, San Francisco, CA 94158, USA; Bruker Nano Surfaces, San Jose, CA, 95134 USA; Institute for Neurodegenerative Diseases, Weill Institute for Neurosciences, University of California, San Francisco, CA, 94158 USA; Amyris Inc., 5885 Hollis St #100, Emeryville, CA, 94608 USA; Department of Chemistry, University of Michigan, Ann Arbor, MI 48104, USA; Chan Zuckerberg Biohub, San Francisco, CA 94158, USA

**Keywords:** microbowls, microwell arrays, mass spectrometry imaging

## Abstract

Patterned surfaces can enhance the sensitivity of laser desorption ionization mass spectrometry by segregating and concentrating analytes, but their fabrication can be challenging. Here, we describe a simple method to fabricate substrates patterned with micron-scale wells that yield more accurate and sensitive mass spectrometry measurements compared to flat surfaces. The wells can also concentrate and localize cells and beads for cell-based assays.

## 1. Introduction

Matrix assisted laser desorption ionization (MALDI) is a form of soft-ionization mass spectrometry (MS) commonly used in biological research for proteomics and metabolomics^[1– 3]^. The ability to rapidly process multiple samples in parallel without auto-feeders makes it suited to high-throughput and single-cell applications^[4–6]^. Key to the method are the matrices or engineered substrates that facilitate the generation of ionic species using energy from a laser ^[7,8]^. The properties of the substrates, including their chemistry, conductivity, and micropatterning impact sample ionization efficiency and, thus, measurement sensitivity^[8–11]^. For example, micron-scale wells are useful for segregating samples of distinct composition so they can be separately analyzed^[12–14]^. Well arrays are also compatible with active^[15,16]^ or passive loading techniques^[12,17]^, to simplify preparation of samples for analysis. However, MALDI-MS requires samples be dried prior to analysis. When droplets are dried on flat surfaces, they tend to distribute their analytes about the perimeter due to the coffee ring effect^[18,19]^. Similar processes occur in cylindrical wells, leading to precipitation along the periphery^[20,21]^ where signal is inhibited due to laser occlusion by the walls. The result in both cases is lowered sensitivity and increased measurement variability due to inhomogeneity of the sample spots ^[18,22]^.

Bowl-shaped wells with curved bases are advantageous because, upon drying, precipitated analytes concentrate at the center in a more uniform fashion^[23]^, where they are efficiently ionized^[24]^. These wells, however, are difficult to fabricate, requiring micromachining, subtractive etching methods, embossing of plastics or specialized deposition methods^[23,25–28]^. Additive fabrication approaches are superior because they use photolithographic techniques that are simple, inexpensive, and ubiquitous. However, photolithographic fabrication of wells with controlled curvature is challenging, requiring complex mask and lens systems to modulate light intensity with micron resolution across the array^[29,30]^. Consequently, simpler but inferior methods with stamping or backfilling of sharp features with polymer are more common, even though they are tedious and provide limited control^[31–34]^. A superior approach would use simple photolithographic techniques without sacrificing control over well shape and curvature.

In this paper, we describe a simple method to photolithographically fabricate wells with controlled curvature in SU8. This photoresist has several properties useful for mass spectrometry, including chemical robustness, precision control of surface features, and scalable fabrication. Using the approach, we fabricate curved-bottom wells at a density of 100,000 per square centimeter on a glass slide. To demonstrate the utility of these wells, we show they yield enhanced sensitivity in microscale mass spectrometry compared to cylindrical wells. Additionally, we show that the curved profiles generate a gravity well that efficiently aggregates cells. The simplicity and control of our approach should make it valuable for single and multi-cell MALDI-MS and for aggregating cells in organoid, embryoid, and cell-cell interaction studies.

## 2. Results and discussion

### 2.1 Microbowls with controlled curvature

SU-8 is composed of bisphenol epoxy resin and the dissolved salt of the photoinitiator triarylsulfonium/hexafluoroantimonate. Upon exposure to ultraviolet (UV) light (**Figure 1a**), the photoinitiator degrades into hexafluoroantimonic acid, which protonates the bisphenol epoxides such that, when heated, they crosslink into a durable polymeric material^[35,36]^. The final structure of the crosslinked network thus depends on the concentration of activated hexafluoroantimonic acid in the SU-8. Since this acid is a dissolved small molecule, it can diffuse post-exposure into unexposed regions, crosslinking them, and thereby allowing transformation of wells with vertical walls and flat bases into ones with sloped walls and curved bases (**Figure 1b**). Because the final shape depends on the diffusion profile of the acid, it can be controlled by the time and temperature over which the substrate is incubated prior to solvent development, which removes the uncrosslinked SU-8 and halts fabrication (**Figure 2a** and **b**).

**Figure 1.**
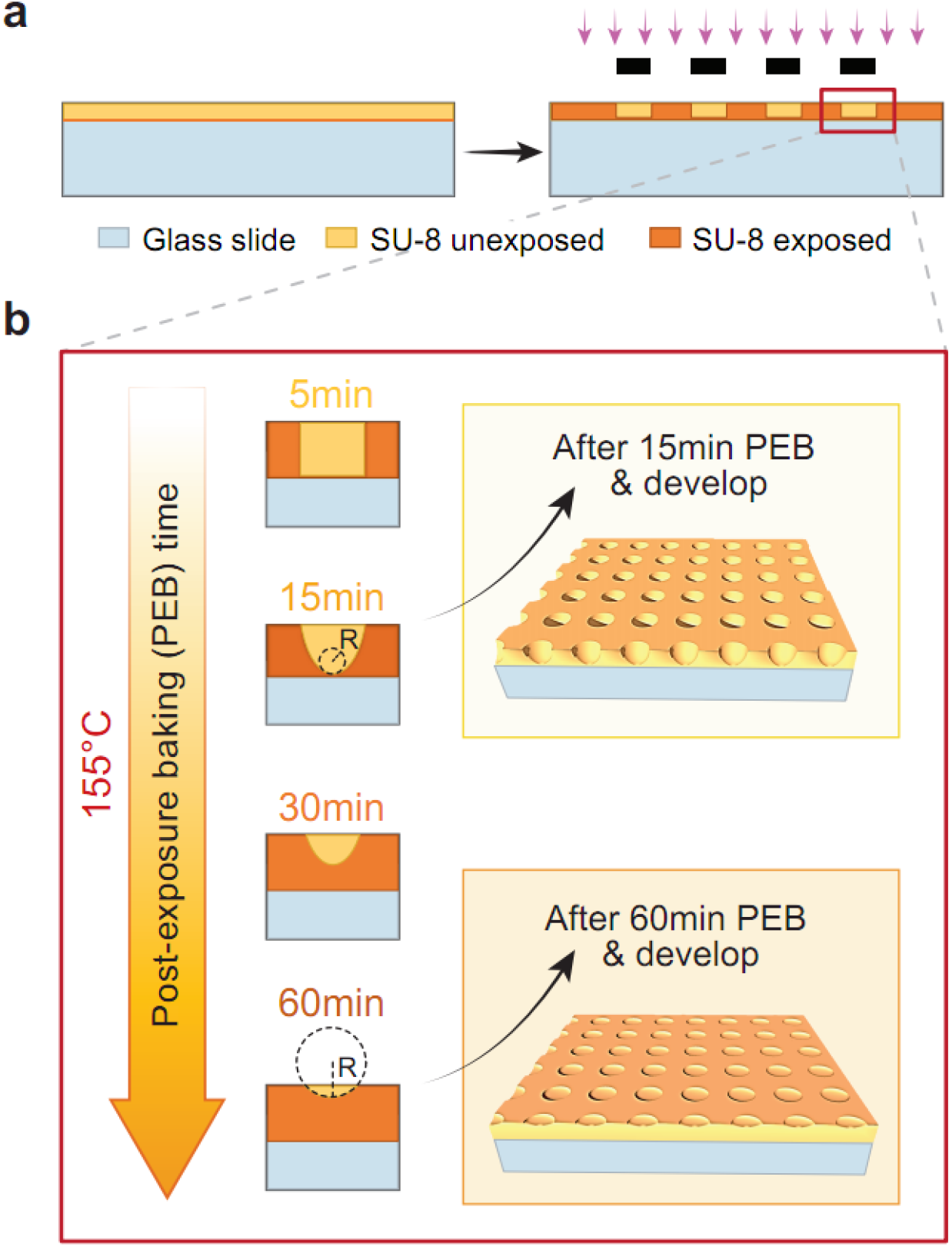
Overview of the fabrication process of microwells. a. Spin coat SU 8 on top of a glass slide and then transfer patterns of microwells from the mask to SU 8 by UV exposure. b. By controlling the PEB time at escalated temperature, the curvature of the profile of the microwells can be adjusted accordingly.

**Figure 2.**
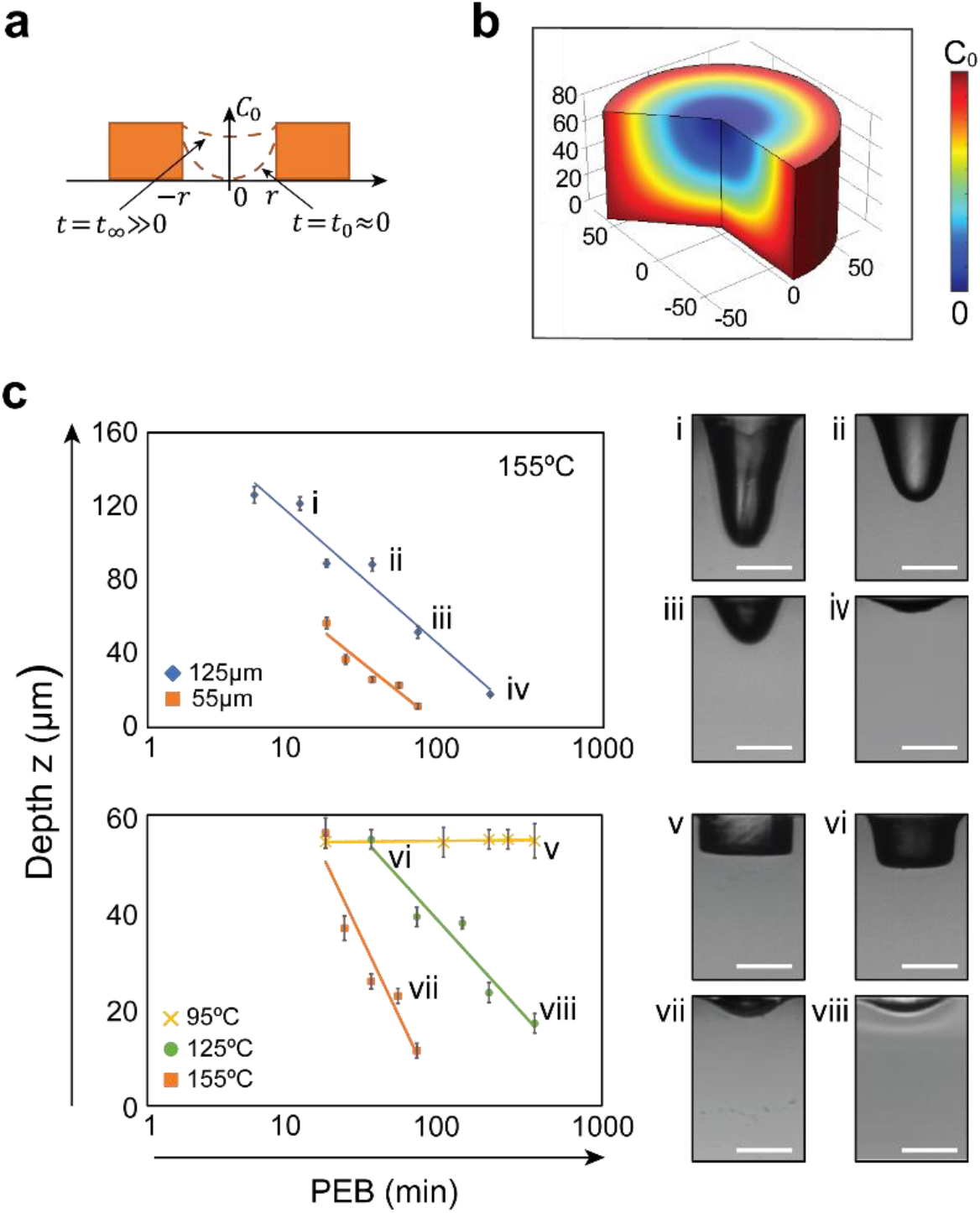
Prediction and validation of relationships between the profile of the microwells with PEB time. **a**. Profile of the microwell changes with PEB time. At time zero, few photo initiators diffuse into the unexposed microwell and the profile of the microwell is still close to cylinder shape. At time infinity, as the concentration of photo initiators in the unexposed microwell reaches the exposed part of the SU 8, *C*_0_, the profile of the microwell will be a shallow curve with large curvature. **b**. COMSOL simulation result based on Fick’s diffusion law. **c**. Experiment validation of the proposed theory based on the diffusion of photo initiators. Top graph shows the depth of the microwell versus PEB time when PEB temperature is set to be 155 °C. Blue diamonds and orange squares are measurements from the microwells when the SU 8 layer is 125 µm and 55 µm respectively. Bottom graph shows how the depth of the microwell changes with PEB time when the thickness of SU 8 layer is fixed to 55 µm. Yellow cross, green dots, and orange squares are measurements from the microwells at different PEB temperature 95°C, 125°C and 155°C respectively. Error bars show the standard deviation of five measurements. All scale bars are 50 µm and fit lines natural logarithms.

To investigate this, we vary the time and temperature of the post-exposure bake (PEB) and thickness of the SU-8 layer and measure the resultant well profiles. When we fix temperature at 155 °C and vary the thickness of the SU-8 layer, the well depth follows a log-time relationship (**Figure 2c**, upper). Alternatively, when we fix thickness to 55 µm and change temperature, the slope changes, indicating that the SU-8 liquifies above a threshold temperature. At 95°C (yellow crosses and fit line), the wells do not change appreciably over the PEB, indicating that the SU-8 remains solid and diffusion of the photoacid is minimal; this is the PEB temperature recommended by the manufacturer to maintain sharp features. By contrast, when we increase PEB temperature to 125°C (green circles and fit line) or 155°C, the wells become rounded, with the change occurring faster at higher temperature than at lower (**Figure 2c**, lower, orange and green). Moreover, because microbowl depth depends on temperature and PEB time, which can be finely tuned and kept constant across the substrate, the microbowls are uniform in shape and depth (**Supplementary Figure 1**).

Because the drying pattern of samples can influence the signal sensitivity and variation of MALDI MS, we investigate how the morphology of the surface and the shape of the microwells influences drying and the resultant analyte deposition. Using an inkjet printer, we print arrays of water droplets containing 5 µm fluorescent polystyrene beads (0.05% w/v) onto an SU-8 substrate patterned with microbowls, cylindrical microwells, or as a flat surface (Methods, **Figure 3a**). The 500 pL droplets dry within 10 mins at room temperature, leaving behind the beads. To characterize the drying behavior, we measure the final displacement of the beads from the printed droplet’s center (**Figure 3b, Supplementary Figure 2**). For the flat surface, the beads tend to deposit at the periphery of the original droplets likely due to the coffee ring effect (**Figure 3a iii, Video S1**). For cylindrical wells, the beads tend to also deposit at the periphery (**Figure 3a ii, Video S2**). By contrast, for microbowls, the beads tend to deposit at the center, indicating suppression of the coffee ring effect^[37]^ (**Figure 3a i, Video S3**). This is beneficial because deposition at the periphery tends to dilute analyte concentration, reducing signal intensity and interfering with reproducible laser scanning since the precipitated analyte position varies from well to well^[24]^. By contrast, with microbowls, the analytes are always at the center, and maximally concentrated.

**Figure 3.**
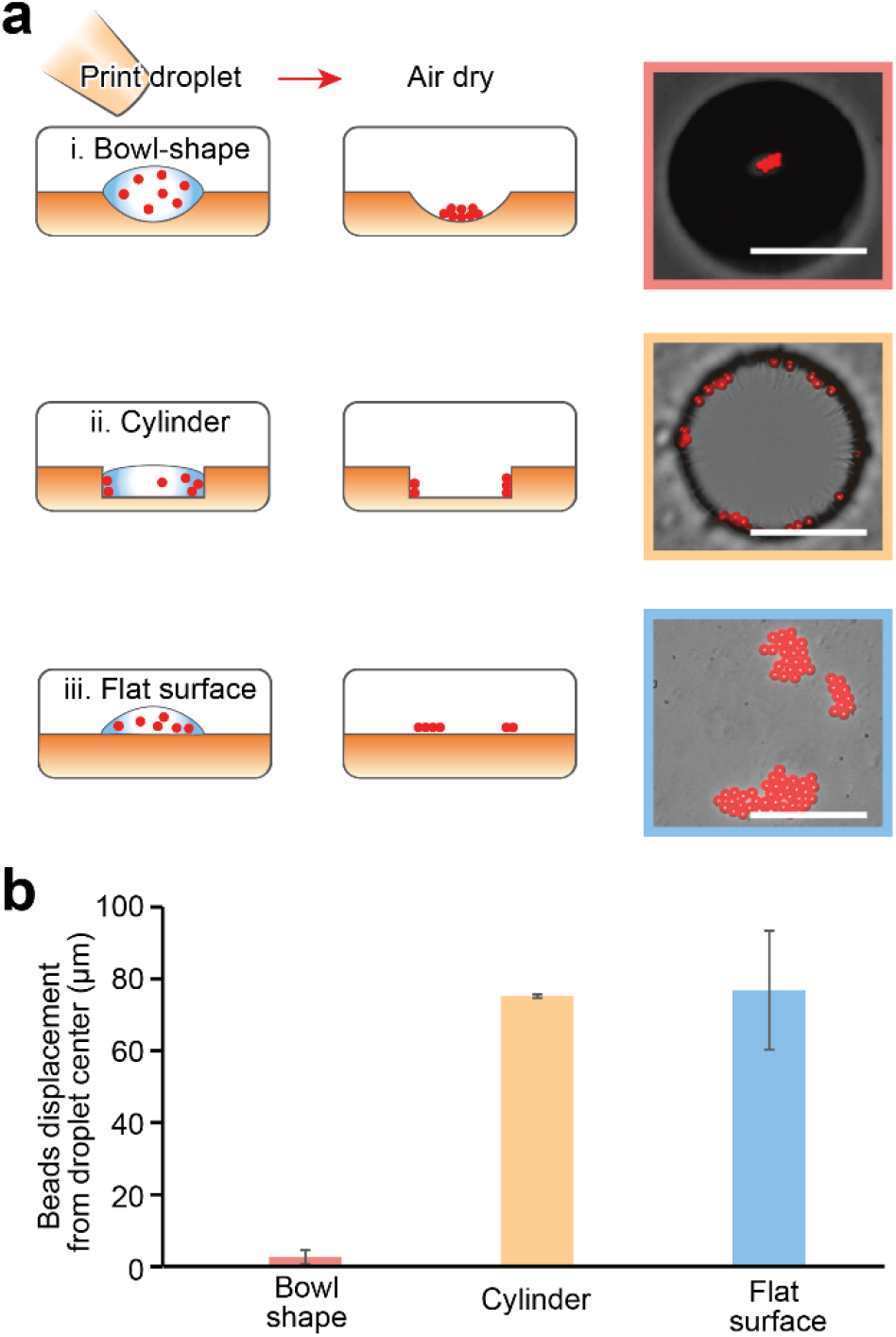
Drying patterns of microbeads on substrates with different morphology. **a**, Droplets containing microbeads are inkjet printed on three types of substrate: **i**, microbowls, **ii**, cylindrical wells and **iii**, flat surface from left top to bottom. Right panels show the distribution of microbeads after drying. **b**, Average displacement of microbeads after drying to the center of the droplet original printed droplet. Scale bars are 100 µm. Error bars are standard deviations of ten measurements.

An important application of wells with controlled curvature is microscale mass spectrometry (µMS), a technique allowing high-density analysis of thousands of samples in parallel. In the technique, samples are confined in wells where they can be energized with the ionizing laser of MALDI. The curvature of the wells is important because MALDI requires that the samples be desiccated prior to deposition of matrix. Microbowls tend to precipitate analytes at the center, localizing and concentrating them in a laser-accessible region. The result is higher sensitivity and reduced measurement error compared to cylindrical wells. To illustrate this, we fabricate ten thousand microbowls above a conductive glass substrate. To load the wells, we use a commercial droplet dispenser for continuous reagents or printed droplet microfluidics (PDM) for suspensions, which overcomes Poisson loading (Methods), although other systems capable of picoliter dispensing are applicable^[38,39]^. We fabricate the wells with a depth of 15.1 µm ±2.3 µm µm (**Supplementary Figure 1**) to allow MALDI-MS analysis which has specific requirements on sample depth (**Methods**). We load the wells with rows of droplets (100 µm in diameter) containing 0, 10, 100 and 500 ng/mL naringenin in DI water, respectively (**Figure 4a**). After printing, we place the sample in a desiccator to dry (**Figure 4b**). We mount the dried substrate onto the requisite MALDI-MS imaging adapter, coat with matrix, and analyze with the instrument (**Figure 4b, c** and **d, Methods**). For comparison, we repeat this process for the flat surfaces and cylindrical wells (**Figure 4c** and **d**). Comparison of the substrates in fluorescence mode shows that while microbowls precipitate analytes at the center and distribute matrix evenly across the whole well, cylindrical wells distribute them on the perimeter and matrix forms random patterns on the flat surface (**Figures 4b** and **c**). We determine the limits of quantification (LoQ)^[40]^ of naringenin are 34.98 ng/mL, 55.59 ng/mL and 228.79 ng/mL for microbowls, flat surface and cylindrical wells, respectively (**Figure 4e, Supplementary Figure 3** and **Method**). Because the analyte is distributed over a larger area in cylindrical wells, it is diluted while also being less accessible to the matrix and laser, resulting in lowered and noisier signals compared to microbowls. For the flat surface, due to the coffee ring effect, analytes also distribute unevenly. In addition, matrix tends to dry unevenly due to the lack of pinning features, resulting in significantly noisier and reduced signals compared to microbowls (**Figure 4e, Supplementary Figure 3**). To characterize the impact of microbowl depth, we repeat these experiments with different depth (15-µm and 50- µm) and find that shallower wells yield enhanced signal (**Supplementary Figure 4**). This is likely due to the dependence of the MALDI signal on sample location, which must be matched with the focal plane of the ionizing laser (**Methods**).

**Figure 4.**
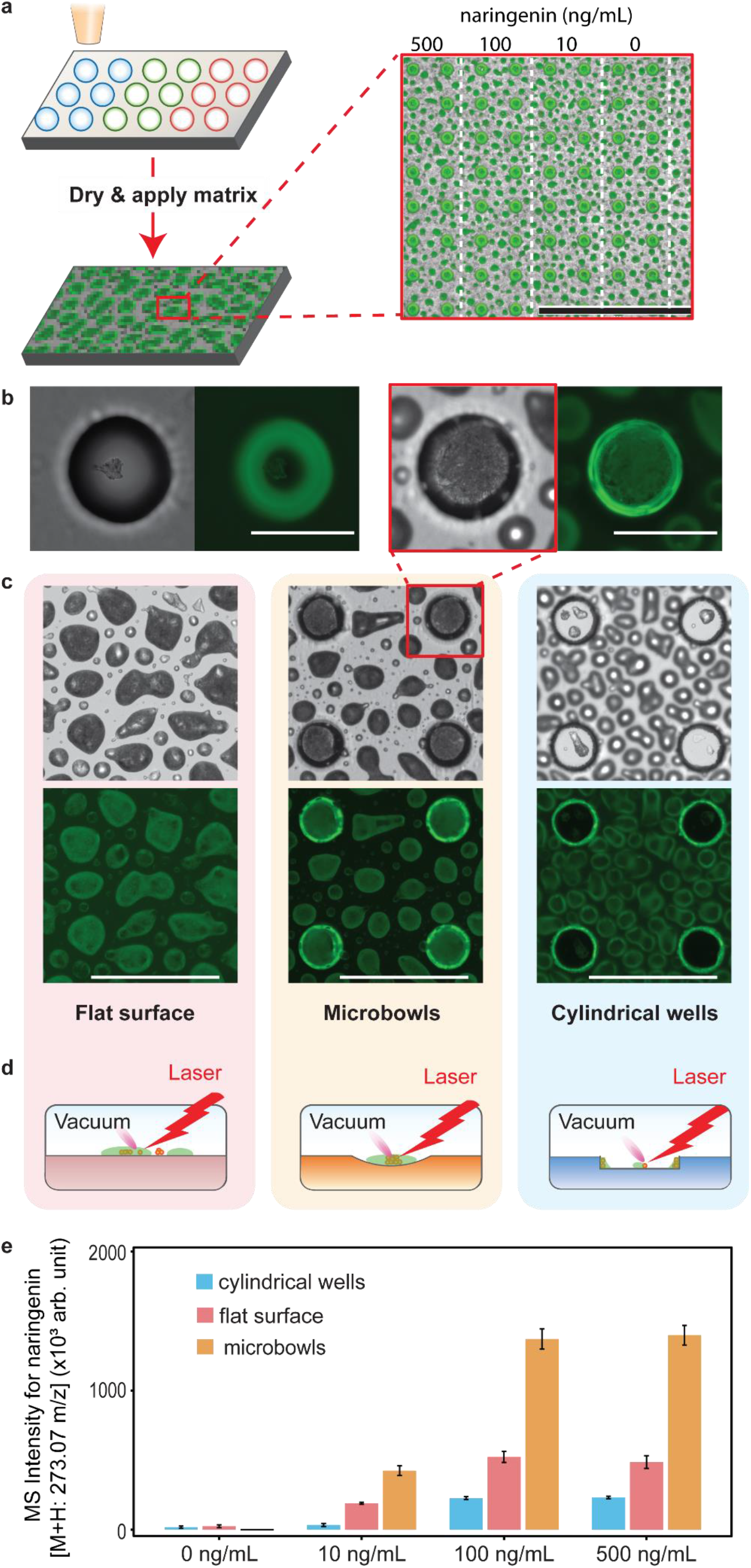
Microbowls concentrate analytes, enhancing MS signal. **a**, Droplets containing different concentrations of naringenin are printed on substrates with different morphology. After the droplets are dried at room temperature, matrix solutions are applied. Inset shows an array of microbowls printed with different concentrations of naringenin. Scale bar is 1 mm. **b**, Combined image of transillumination and GFP epifluorescence channels of dried droplet in a microbowl before (left two images) and after the coating of matrix (right two images). Scale bars are 100 µm. **c** Printed substrates after spray-coating of matrix, with transillumination and GFP epifluorescence images of the flat surface (light red), microbowls (light yellow) and cylindrical wells (light blue). Scale bars are 300 µm. **d**, Illustrations of ionizing laser and substrate geometry. **e**, Sums of naringenin peak amplitude in MS signal for flat surface, microbowls, and cylindrical wells. Error bars show the standard deviation of six samples.

To determine whether these properties extend to the detection of other molecules, we use microbowls to quantify triacetic acid lactone (TAL) and two small peptides and observe similar enhancements (**Supplementary Figure 5, Methods**). Thus, due to their even distribution of matrix and analyte concentrating power, microbowls yield increased and more uniform signals than other common substrate geometries.

Mass spectrometry allows label-free quantitation of a broad range of analytes, making it ideal for characterizing samples of unknown composition. An especially important area in which this is useful is the analysis of microbes engineered to express exogenous metabolisms, including for biocircuitry and bioproduction. Libraries of microbes engineered to express millions of exogenous pathways can be efficiently generated with genetic methods, but testing each variant for the phenotype of interest is laborious, requiring isolation, cultivation, and analysis in separate wells of microtiter plates. µMS is a significant advance because it allows parallel analysis of tens of thousands of variants per square centimeter of substrate, reducing the scanning time and volume of reagents consumed. Key to such screens is accurately characterizing the metabolisms of the variants, which requires an accurate and sensitive measurement of metabolites. Because microbowls enhance µMS signals, they are an invaluable feature of such microbial screens. To illustrate this, we use the approach to analyze two yeast strains engineered to produced naringenin, a chemical that has been shown to have a range of potential therapeutic uses ^[41]^, low producer and high producer strains. Because a single yeast cell does not produce sufficient material for µMS analysis, we pre-culture the strains in microfluidic droplets, generating colonies comprising thousands of genetically identical cells (**Figure 5a**, Steps 1 and 2). To induce production of naringenin, introducing medium is added to the culture droplets by merger, which has the inducer required to activate compound production (**Method**). The colonies are cultured for another week before being dispensed to the microbowls using PDM. PDM ensures that every well is loaded with a colony and to pattern them on the substrate so that we can directly compare the production of naringenin for the two strains (**Figure 5b**). The substrate is then processed through µMS using the standard workflow, removing the printing oil, drying the samples in a desiccator, applying the matrix, and scanning (**Figure 5a** Step 5, and **5c**). By eye there is a clear difference between the two strains according to the printing grid (**Figure 5c**). To quantify this difference, we measure the intensity distribution for the centers of the wells, finding that, indeed, the high producer is more efficient at making this molecule than the low producer (**Figure 5d**). Using recently described approaches, cells with desired properties can be recovered from the array and sequenced^[12]^. Additionally, we demonstrate differentiation of yeast strains engineered to produce varying levels of TAL (**Supplementary Figure 6**).These results show that µMS with microbowls can sensitively quantify the bioproduction of exogenous molecules in a label-free fashion appropriate for high throughput screening.

**Figure 5.**
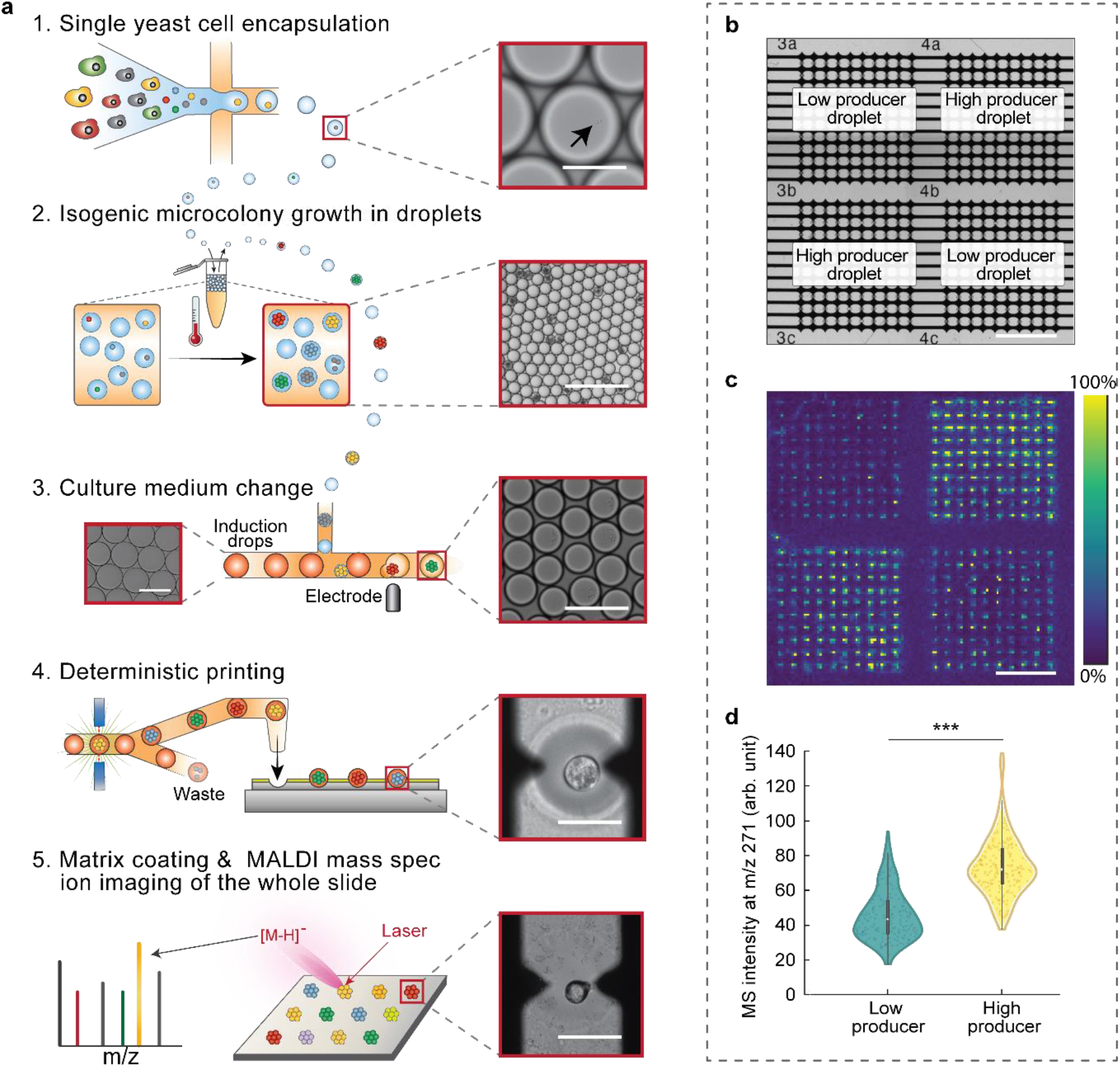
Microscale mass spectrometry measurement of two yeast strains engineered to produce different amounts of naringenin. The two strains were prepared and printed separately using the same protocol. **a**, Step 1, genes of interest are integrated into yeast, which are individually encapsulated in droplets containing culture medium. Inset shows a droplet containing a single yeast cell. Scale bar is 40 µm. Step 2, droplets are incubated to form isogenic microcolonies. Inset shows droplets containing the yeast microcolonies. Scale bar is 200 µm. Step 3, after incubation, each droplet is merged with another larger droplet containing culture medium that is suitable for Naringenin production (pathway expression inducing medium). Left inset shows the media droplets with scale bar 100 µm. Right inset shows the merged droplets with scale bar 200 µm. Step 4, droplets containing isogenic microcolonies of yeast are printed into microbowls at defined positions. Inset shows a droplet within a microbowl immediately after printing, with scale bar equals 100 µm. Step 5, after matrix is sprayed, MALDI MS ion imaging is performed. Inset shows a dried droplet containing a microcolony in a microwell, with scale bar 100 µm. **b**, Bright-field transillumination image showing layout of printed strains within the microbowl array. Scale bar is 1 mm. **c**, MALDI MS ion image shows the MS signal for naringenin. Scale bar is 1 mm. **d**, Violin plot compares the MS signal of Naringenin from microwells printed with high producers and low producers, respectively. Error bars show the standard deviation of 200 samples. ***Populations are significantly different (*p* < 0.001).

### 2.2 Microbowls efficiently aggregate cells for interaction studies

Interaction studies are essential for characterizing cooperative phenotypes between cells, which are important in applications like T cell killing of cancer cells and cultivation of wild microbes. Performing such studies requires that the interacting cells be brought together and monitored. Common cylindrical wells are poorly suited to this, because the flat bottoms do not force cells together, while the vertical walls occlude visualization near the edges. Microbowls afford an effective alternative, providing a gravity well that draws cells to the center where they are visible. To demonstrate the utility of microwells for cell interaction studies, we use them to observe interactions between killer T cells (NK92) and cancer cells (K562). We load the cells in the wells randomly by sedimentation at an average of one cell per well, yielding ∼36% of wells containing one of each cell type. For comparison, we repeat the process with cylindrical wells. As expected, we find that cells in microbowls reliably contact at the centers, where they interact and are easily imaged (**Figure 6c** and **d**, NK92 cells are labeled green and K562 cells are labeled red). By contrast, with cylindrical wells, the cells are randomly distributed and do not reliably interact **(Figure 6f** and **g**). Consequently, while we observe robust killing of K562 cells in microbowls, we do not in cylinders (**Figure 6e and 6h**). These results demonstrate that microbowls ensure cell contact in interaction assays, which can impact observations. After culturing cells in microbowls, we also have tested single cell MS using microbowls (**Supplementary figure 7**). However, in order to keep the right osmosis pressure for the cells during droplet printing, we have to use PBS solution to encapsulate the single cell but the high salt concentration produces a significant background noise, which prevents the detection of the signals from the single cell. We expect that with the built-in HPLC function in the DESI MS instrument, it should be solved in the future.

**Figure 6.**
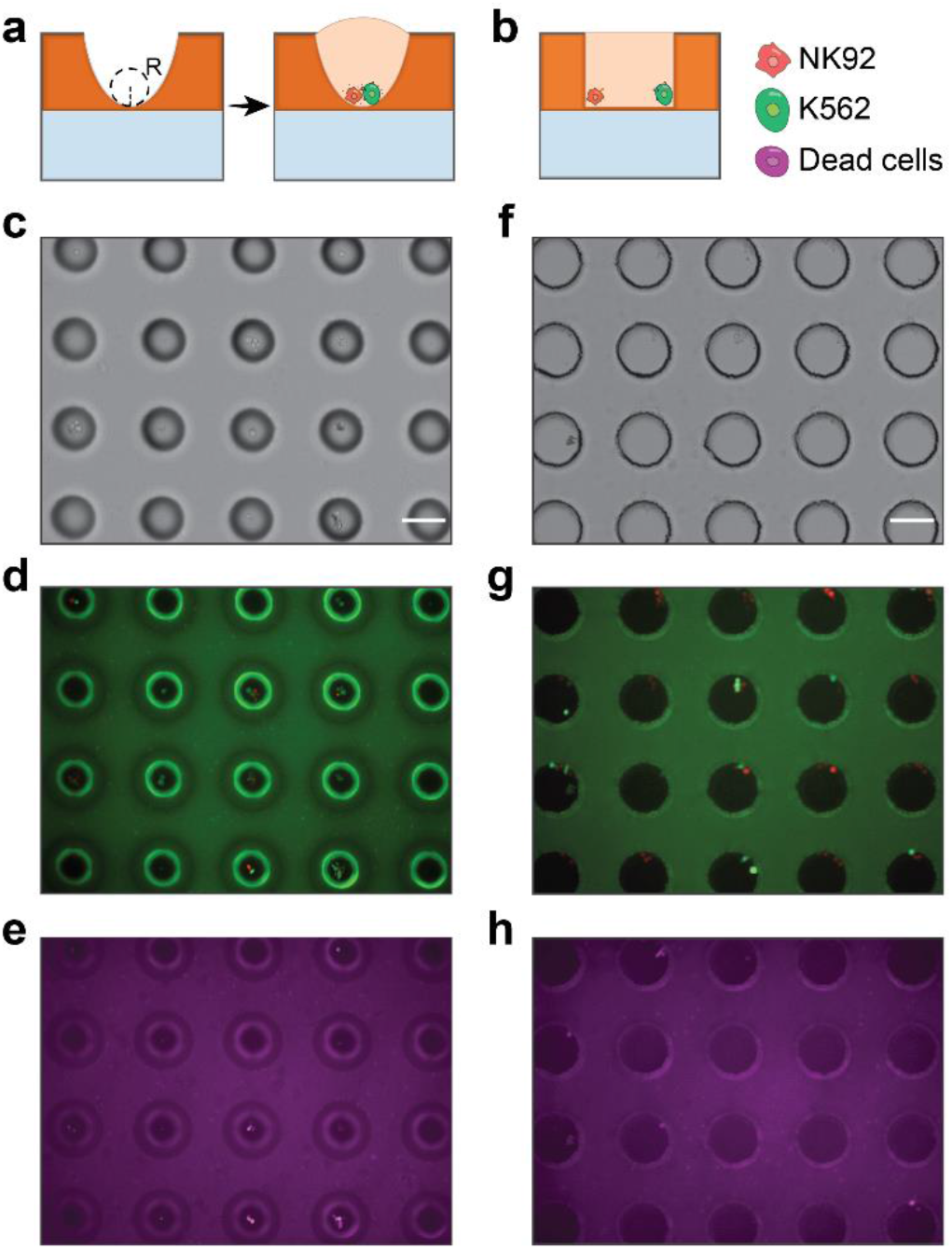
Cell interaction study of microbowls with a profile of deep and sharp radius of curvature at the bottom. a. Due to the microwells are deep and has a sharp radius of curvature at the bottom, they are easy to trap cells by passive loading and pull the cells in close contact to facilitate cell interactions. While for b, cylindrical wells, cells tend to distribute randomly inside and have less chance to get in touch with each other. c, d and e show the transillumination image, combine of GFP and RFP epifluorescence image and Cy5 epifluorescence image of the microbowls respectively. Similarly, f, g and h show the images of cylindrical wells. All scale bars are 100 µm.

## 3. Conclusions

We describe a simple and controlled method to fabricate microbowl arrays. Our approach generates bowls with a range of shapes, sizes, and vertical profiles by controlling post-exposure bake time and temperature. The microbowls are useful for numerous applications, especially for enhancing the sensitivity of microscale mass spectrometry. In addition, the wells can be loaded via active printing or passive Laplace guidance techniques^[15,17,38]^, automating preparation of thousands of samples for analysis. The planar grid is compatible with spatial indexing to allow integration of imaging, µMS, and sequencing.^[12,42]^ The sensitivity enhancement of µMS achieved with microbowls is useful for characterizing engineered metabolisms for biocircuits and biomolecule production. These screens can be used to optimize pathways or enzymes to enhance production of target analytes, or detect novel products through metabolic biosensing^[43,44]^. In addition, as we have shown, they provide a gravity well by which to bring cells into contact to ensure interaction, which is important for functional screens of B and T cells that comprise cell therapies. When combined with printed droplet microfluidics, every well can be loaded with an exact number of different cell types, allowing tens of thousands of interaction studies per square centimeter of slide. Similar methods are useful for aggregating cells of different types into seeds that can form spheroids, organoids, and embryoids, but with higher efficiency and control compared to existing techniques. Thus, while simple, microbowls provide a valuable platform for studies involving the functional and multiomic analysis of single cells and multi-cell consortia.

## Supporting Information

Supporting Information is available from the Wiley Online Library.

## Competing Interests

The authors declare no competing interests.

## Author Contributions

L.X. conceptualized the project anddesigned the experiments. X.L. prepared the mamaliam cells. W. L., J. H., C. L. prepared the yeast cells. L.X., K. C., P. Z. performed the droplet and cell printing. L. X., N. T., H. Y. performed the MALDI MS imaging, L.X. K. C., E. P., A. A. analyzed and interpreted the obtained data. L. X., C. M., A. A. wrote the manuscript. All authors read and agreed to the final work.

## Acknowledgements

This work was supported by the Chan Zuckerberg Biohub, the National Institutes of Health (NIH) (Grant Nos. R01-EB019453-02 and R01-HG008978-01), Office of the Director of National Intelligence (ODNI) (IARPA FELIX Contract No. N66001-18-C-4507), United States Department of Defense, Defense Advanced Research Projects Agency (DARPA) (Agreement No. W911NF1920013 The content of the information does not necessarily reflect the position or the policy of the Government, and no official endorsement should be inferred.) and funded in part by federal funds from the Virology Surveillance and Diagnosis Branch, Influenza Division, Centers for Disease Control and Prevention, under BAA 75D301-19-R-67835 (Topic #6). We thank Kirsten Benjamin, John Hung, Yue Yang, Matthew Rienzo, Adam Navidi, Michael Leavell for helpful discussion, and Mona Elbadawi and Carol Tran for strain analysis.

## Notes

### Competing Interest Statement

The authors have declared no competing interest.

## References

[1] D. C. Castro, Y. R. Xie, S. S. Rubakhin, E. V. Romanova, J. V. Sweedler, Nat Methods 2021, 18, 1233.

[2] R. Kaufmann, J Biotechnol 1995, 41, 155.

[3] M. Kompauer, S. Heiles, B. Spengler, Nat Methods 2017, 14, 90.

[4] Z. Yuan, Q. Zhou, L. Cai, L. Pan, W. Sun, S. Qumu, S. Yu, J. Feng, H. Zhao, Y. Zheng, M. Shi, S. Li, Y. Chen, X. Zhang, M. Q. Zhang, Nat Methods 2021, 18, 1223.

[5] A.J. Ibáñez, S. R. Fagerer, A. M. Schmidt, P. L. Urban, K. Jefimovs, P. Geiger, R. Dechant, M. Heinemann, R. Zenobi, Proc Natl Acad Sci U S A 2013, 110, 8790.

[6] T. Si, B. Li, T. J. Comi, Y. Wu, P. Hu, Y. Wu, Y. Min, D. A. Mitchell, H. Zhao, J. V. Sweedler, J Am Chem Soc 2017, 139, 12466.

[7] S. M. A. B. Batoy, E. Akhmetova, S. Miladinovic, J. Smeal, C. L. Wilkins, Applied Spectroscopy Reviews 2008, 43, 485.

[8] H. Togashi, Y. Kobayashi, Rapid Commun Mass Spectrom 2009, 23, 2952.

[9] M. Benz, A. Asperger, M. Hamester, A. Welle, S. Heissler, P. A. Levkin, Nat Commun 2020, 11, 5391.

[10] J. Bai, Y. H. Liu, D. M. Lubman, D. Siemieniak, Rapid Commun Mass Spectrom 1994, 8, 687.

[11] F. Fournelle, E. Yang, M. Dufresne, P. Chaurand, Anal Chem 2020, 92, 5158.

[12] J. C. Love, J. L. Ronan, G. M. Grotenbreg, A. G. van der Veen, H. L. Ploegh, Nat Biotechnol 2006, 24, 703.

[13] F. Khan, R. Zhang, A. Unciti-Broceta, J.J. Díaz-Mochón, M. Bradley, Advanced Materials 2007, 19, 3524.

[14] M.-H. Kang, J. Park, S. Kang, S. Jeon, M. Lee, J.-Y. Shim, J. Lee, T. J. Jeon, M. K. Ahn, S. M. Lee, O. Kwon, B. H. Kim, J. R. Meyerson, M. J. Lee, K.-I. Lim, S.-H. Roh, W. C. Lee, J. Park, Advanced Materials n.d., n/a, 2102991.

[15] R. H. Cole, S.-Y. Tang, C. A. Siltanen, P. Shahi, J. Q. Zhang, S. Poust, Z. J. Gartner, A. R. Abate, PNAS 2017, 114, 8728.

[16] P. Zhang, A. R. Abate, Advanced Materials 2020, 32, 2005346.

[17] A. Kulesa, J. Kehe, J. E. Hurtado, P. Tawde, P. C. Blainey, PNAS 2018, 115, 6685.

[18] J.-B. Hu, Y.-C. Chen, P. L. Urban, Anal Chim Acta 2013, 766, 77.

[19] Y.-H. Lai, Y.-H. Cai, H. Lee, Y.-M. Ou, C.-H. Hsiao, C.-W. Tsao, H.-T. Chang, Y.-S. Wang, J Am Soc Mass Spectrom 2016, 27, 1314.

[20] B. Rieger, L. R. van den Doel, L. J. van Vliet, Phys. Rev. E 2003, 68, 036312.

[21] W. Sun, F. Yang, Langmuir 2015, 31, 4024.

[22] G. McCombie, R. Knochenmuss, J Am Soc Mass Spectrom 2006, 17, 737.

[23] W. Li, M. Khan, H. Li, L. Lin, S. Mao, J.-M. Lin, Chem Commun (Camb) 2019, 55, 2166.

[24] O. Kudina, B. Eral, F. Mugele, Anal. Chem. 2016, 88, 4669.

[25] Y. Xu, F. Xie, T. Qiu, L. Xie, W. Xing, J. Cheng, Biomicrofluidics 2012, 6, 016504.

[26] M. Nikkhah, J. S. Strobl, V. Srinivasaraghavan, M. Agah, IEEE Sensors Journal 2013, 13, 1125.

[27] T. Liu, C.-C. Chien, L. Parkinson, B. Thierry, ACS Appl Mater Interfaces 2014, 6, 8090.

[28] Z. Li, X. Guo, L. Sun, J. Xu, W. Liu, T. Li, J. Wang, Biotechnol Bioeng 2020, 117, 1092.

[29] L. Zhou, X.-X. Dong, G.-C. Lv, J. Chen, S. Shen, Optics Communications 2015, 342, 167.

[30] K. Zhong, Y. Gao, F. Li, Z. Zhang, N. Luo, Optik 2014, 125, 2413.

[31] E. J. Vrij, S. Espinoza, M. Heilig, A. Kolew, M. Schneider, C. A. van Blitterswijk, R. K. Truckenmüller, N. C. Rivron, Lab Chip 2016, 16, 734.

[32] T. Si, B. Li, T. J. Comi, Y. Wu, P. Hu, Y. Wu, Y. Min, D. A. Mitchell, H. Zhao, J. V. Sweedler, J. Am. Chem. Soc. 2017, 139, 12466.

[33] J. Y. Park, C. M. Hwang, S.-H. Lee, Biomed Microdevices 2009, 11, 129.

[34] G. S. Jeong, J. H. Song, A. R. Kang, Y. Jun, J. H. Kim, J. Y. Chang, S.-H. Lee, Advanced Healthcare Materials 2013, 2, 119.

[35] H. Lorenz, M. Despont, N. Fahrni, N. LaBianca, P. Renaud, P. Vettiger, J. Micromech. Microeng. 1997, 7, 121.

[36] K. V. Nemani, K. L. Moodie, J. B. Brennick, A. Su, B. Gimi, Materials Science and Engineering: C 2013, 33, 4453.

[37] P. J. Yunker, T. Still, M. A. Lohr, A. G. Yodh, Nature 2011, 476, 308.

[38] M. Singh, H. M. Haverinen, P. Dhagat, G. E. Jabbour, Advanced Materials 2010, 22, 673.

[39] A. Ovsianikov, M. Gruene, M. Pflaum, L. Koch, F. Maiorana, M. Wilhelmi, A. Haverich, B. Chichkov, Biofabrication 2010, 2, 014104.

[40] D. A. Armbruster, T. Pry, Clin Biochem Rev 2008, 29, S49.

[41] C. M. Palmer, K. K. Miller, A. Nguyen, H. S. Alper, Metabolic Engineering 2020, 57, 174.

[42] A.J. Ibáñez, S. R. Fagerer, A. M. Schmidt, P. L. Urban, K. Jefimovs, P. Geiger, R. Dechant, M. Heinemann, R. Zenobi, PNAS 2013, 110, 8790.

[43] J. Nielsen, Appl Microbiol Biotechnol 2001, 55, 263.

[44] L. Xu, K.-C. Chang, E. M. Payne, C. Modavi, L. Liu, C. M. Palmer, N. Tao, H. S. Alper, R. T. Kennedy, D. S. Cornett, A. R. Abate, Nat Commun 2021, 12, 6803.

