## Supplementary material for "Microbowls with controlled concavity for accurate microscale mass spectrometry": Method and Supplementary figures

### MATERIALS AND METHODS

#### Mammalian cell culture and staining

Three cell lines were used in the experiment: 3T3 (ATCC CRL-1658) K562 (ATCC CCL-243) and NK-92® MI (ATCC CRL-2407). Cells were cultured under the supplier's recommended conditions. The NK-92® MI were cultured in MyeloCult™ H5100 (StemCell Technologies cat # 05150) at 37 °C under 5% $CO_2$. The 3T3 and K562 were cultured in Iscove's Modified Dulbecco's Medium (ATCC 30-2005) with 10% fetal bovine serum (Gibco 10437028) at 37 °C under 5% $CO_2$ respectively. NK-92® MI cells and K562 cells were stained with Calcein Green AM (ThermoFisher C34852) and Calcein Red-Orange AM (ThermoFisher C34851), respectively, prior to loading into the microbowls arrays for cell-cell interaction experiment. 7-Aminoactinomycin D (7-AAD, ThermoFisher A1310) was used to stain dead cells. Similarly, for single cell MS analysis, 3T3 and K562 cells are cultured and then stained with CellTracker Red CMTPX (ThermoFisher C34552) and CellTracker green CMFDA (ThermoFisher C2925) in PBS (ThermoFisher 14040141) respectively. After staining, the cells are ready for droplet encapsulation and droplet printing into the microbowls arrays for MALDI MS.

#### Yeast strain building and culturing

Chemicals were purchased from Thermo Fisher Scientific unless indicated otherwise. All media components were filter sterilized, with the exception of yeast extract, Bacto Peptone, and agar, which were autoclaved before addition of other components. Yeast growth media was composed of a chemically defined basal media called Bird seed medium (BSM) was used in the droplet culture and contained $KH_2PO_4$ (8 g $L^{-1}$ ), $(NH_3)_2SO_4$ (15 g $L^{-1}$ ), $MgSO_4$ (25 mM), succinic acid (50 mM), EDTA (400 μM), $ZnSO_4$ (200 μM), $CuSO_4$ (20 μM), $MnCl_2$ (16 μM), $CoCl_2$ (20 μM), $Na_2^{-1}$ $MoO_4$ (20 μM), $FeSO_4$ (100 μM), $CaCl_2$ (200 μM), biotin (0.6 mg $L^{-1}$ ), p-aminobenzoic acid (2.4 mg $L^{-1}$ ), calcium pantothenate (12 mg $L^{-1}$ ), nicotinic acid (12 mg $L^{-1}$ ), myoinositol (300 mg $L^{-1}$ ), thiamine·HCl (12 mg $L^{-1}$ ), and pyridoxine·HCl (12 mg $L^{-1}$ ), adjusted to pH 5.0 and 2% (20 g/L) sugar comprised of 1.9 % sucrose plus 0.1 % glucose. Yeast pathway expression media was composed of 25% strength BSM (i.e., a four-fold dilution of sugar-free BSM in water) and 4% sugar, comprised of 3.8% galactose and 0.2% glucose. Yeast was cultured in the growth media without galactose for building biomass, and then merged into droplets containing media with galactose to induce naringenin pathway expression.

The *Saccharomyces cerevisiae* yeast strains used in this study were derived from CEN.PK113- 7D [1]. The DNA constructs for yeast transformation and integration into the chromosomes were generated using DNA assembly methods previously described[2,3]. Standard methods were implemented for DNA transformation into competent yeast cells[4,5]. Naringenin-producing strains (both a high-producer and low-producer strain) were designed by an Artificial Intelligence Strain designer nicknamed LILA within Amyris' DARPA-funded M2K program (manuscript in preparation; https://sim.confex.com/sim/2018/meetingapp.cgi/Paper/37552) and constructed using the methods described above. The design involved overexpressing key naringenin pathway proteins, pentafunctional AROM enzyme, DAHP synthase, chorismate mutase, phenylalmine/tyrosine ammonia lyase, trans-cinnamate 4-monooxygenase, 4-courarate-CoA ligase, naringenin chalcone synthase, chalcone isomerase, cytochrome P450 reductase. The high naringenin producer has more integrated gene copies of 4-monooxygenase, 4-courarate-CoA ligase, Naringenin chalcone synthase, chalcone isomerase. All the genes overexpressed for naringenin production were driven by strong galactose-inducible promoters (e.g. pGAL1, pGAL2, pGAL10, pGAL7). In addition, to partially alleviate glucose repression of the GAL promoters, an extra copy of the gene encoding the galactose-responsive transcriptional activator, GAL4, was added with a mutated derivative of its native promoter that lacks the Mig1 repressor binding site (called the GAL4oc* promoter)[6,7]. Strains can be switched from biomass building phase to naringenin production phase by addition of galactose, which triggers the elevated expression of naringenin biosynthetic genes.

The *Yarrowia lipolytica* yeast strains with varied copy numbers of g2ps1 as the high (0.8 mg/mL TAL) and low (0.3 mg/mL TAL) producers for TAL used in this study have been reported[8] and were gifts from Prof. Hal Alper's lab.

All the yeast are first inoculated in a culture tube with 2 mL culture medium separately (Yeast growth media mentioned previously for *Saccharomyces cerevisiae*; 20 g/L glucose, 6.7g/L YNB with ammonium sulfate, and 0.79 g/L CSM in water for *Yarrowia lipolytica*) for one day at 30 °C. The yeasts are then pelleted and resuspended with fresh culture medium. The yeast are counted and diluted with fresh culture medium to a concentration that is ensure roughly 1 in 10 droplets contain a single yeast cell when encapsulated.

**Solution preparation for drying pattern experiments**
5 µm Polystyrene microbeads coated with red fluorescent dye (Spherotech PP-45-10, USA) were diluted in double filtered water at 0.05% w/v ratio. The microbead solution was ultrasonicated at high power (Branson 1510) for 30 mins immediately before use. A reagent spotter (SciFlexArrayer S3, Scienion AG) with a 100 µm capillary nozzle was used to load 300 pL droplets of the prepared microbead solution onto desired positions of the substrate. Videos of different droplets drying were taken from the camera of the spotter.

**Chemical preparation for titration experiments**
For all the titration experimetns, DI water is used as negative control and all the solutions are prepared right before the experiments. Triacetic acid lactone, or TAL (MilliporeSigma H43415), and naringenin (MilliporeSigma 52186) are first dissolved in methanol at 10mg/mL and then the dissolved solutions are further diluted with DI water to 1mg/mL, 500 ng/mL, 100 ng/mL, 10 ng/mL respectively, except for naringenin at 1 mg/mL which does not dissolve totally. Similarly, peptide-mixtures solution with CHCA (MilliporeSigma MSCAL2) at 1µM is diluted with DI water to 500nM, 100nM and 10nM respectively. A reagent spotter (SciFlexArrayer S3, Scienion AG) with a 100 µm capillary nozzle was used to load four 300 pL droplets of the prepared solution onto desired positions of the substrate which are made from ITO slides (MilliporeSigma 50926119).

**Microfluidic devices fabrication**
All the microfluidic devices including the printer head, droplet maker, and droplet merger device were made from poly(dimethylsiloxane) (PDMS) based on the protocols of standard soft lithography[9]. For the printer head, a two-layer SU8 master mold was made. The first SU 8 layer, the flow channel layer for droplet reinjection and sorting, was made to be 80-µm thick and a second layer of 20-µm thick SU 8 was added on top of the first layer only at regions that comprise the channels used to guide insertion of the nozzle and optical fibers. After the master mold was made, uncured PDMS (10:1 polymer to cross-linker ratio) was poured onto the master and cured in an oven at 65 °C for one hour. Then the cured PDMS slab was peeled off before inlet and outlet holes are punched using a 0.75-mm biopsy core. The PDMS casting was then plasma bonded to a 25 mm × 75 mm glass slide and baked at 65 °C overnight. One centimeter of PE/5 tubing (Scientific Commodities) as the printing nozzle of the printer head was inserted into the nozzle channel. The assembled printer head was then treated with AquaPel (AquaPel) and air dried. Similarly, the droplet encapsulation device and droplet merger were made from PDMS with a channel height of 80 µm.

**Yeast encapsulation in droplet, droplet culturing and droplet merging**
Yeast strains with target concentrations in corresponding mediums (Yeast growth media mentioned previously for *Saccharomyces cerevisiae*; 20 g/L glucose, 6.7g/L YNB with ammonium sulfate, and 0.79 g/L CSM in water for *Yarrowia lipolytica*; for the co-culture experiment, 1nM dextran TMR (Catalog # D1818, Thermo Fisher) and 5 nM dextran Cascade blue (Catalog # D1976, ThermoFisher) are added into the culture medium of low and high TAL producer respectively.) and HFE-7500 fluorinated oil (3M) with 2% (w/w) PEG-PFPE amphiphilic block copolymer surfactant (008-Fluoro-surfactant, Ran Technologies) are loaded into separate 1-mL syringes (BD) and injected at 2000 and 7000 µL/h, respectively, into a cross junction droplet maker using syringe pumps (New Era, catalog no. NE-501), controlled with a custom Python script (https://github.com/AbateLab/Pump-Control-Program) (Figure 5a 1). All the droplets are collected into 3-mL syringes (BD) respectively (for the co-culture experiment, droplets of low and high TAL producers are collected in the same syringe). Then the syringes with droplets are fixed on a shaker in a culture room kept at 30 °C. The droplets are incubated in the syringes with a shaking speed at 200 rpm for one week (**Figure 5a 2**). The droplets containing *Yarrowia lipolytica* are ready for next step while the droplets containing *Saccharomyces cerevisiae* need to merge with droplets made of yeast

pathway expression media in a one-to-one fashion: the syringes containing droplets of *Saccharomyces cerevisiae* and yeast pathway expression media (prepared in a cross-junction device as described previously) are injected into a droplet merger, and syringes of HFE-7500 with 2% (w/w) PEG-PFPE amphiphilic block copolymer surfactant are injected as carrying oil phase. Droplet merging is achieved using an electrode connected to a cold cathode fluorescent inverter and DC power supply (Mastech). A voltage of 2.8 V at the power supply produces a ~2 kV AC potential at the electrode, which causes touching droplets to merge (Figure 5a 3).

**Printer head setup and droplet sorting**

A similar optical setup of the printer head from our previous study[10] was used here: a multimode excitation fiber with a core diameter of 105 µm and a Numerical Aperture of 0.10 (Thorlabs) is inserted into a guide channel in the printer head. Similarly, an emission detection fiber with core diameter of 105 µm and Numerical Aperture of 0.22 (Thorlabs) was inserted into a second guide channel in the printer head. Four 50 mW continuous wave lasers with wavelengths of 405, 473, 532, and 640 nm are combined and coupled to the excitation fiber. Emitted light is columnated and ported into a quad-bandpass filter, then passed through a series of dichroic mirrors. Bandpass filters of 448, 510, 571, and 697 nm past each dichroic mirror enable wavelength-specific detection of emitted light by four individual PMTs. Electrode channels and a “Faraday moat” are filled with a 5 M NaCl solution. A positive electrode was connected to a function generator and a high voltage amplifier while a second electrode was grounded. Fluidic inputs into the PDM device are driven by syringe pumps (New Era). Bias and spacer oil containing Novec HFE-7500 oil were flowed through the device at a flow rate of 2000 µL/h. A waste channel was driven with a negative flow rate of − 3000 µL/h. Droplets with yeast cells prepared previously were reinjected into the device at a flow rate of 100 ± 50 µL/h. Real-time optical signal acquisition through a field programmable gate array (FPGA) (National Instruments) was displayed on a customized LabView software. The optical signal was processed in real time and displayed, so droplets of interest (for example, with similar numbers of yeast cells inside) can be identified by specifying gates. Controlled by our custom LabView software, droplets were sorted by passing an oscillating pulse through a high voltage amplifier (Trek 690E-6). Typical droplet sorting parameters range from 10 to 20 kHz, 50 to 100 cycles, and 0.5 to 1.0 kV.

**Well substrate setup**

The well substrate used for PDM is made based on glass slides with chromium electrodes and SU-8 microbowls as described in the main text(**Supplementary figure 1a**). The well substrate was immersed in a bath of Novec HFE7500 oil during printing operation. Copper tape with a conductive adhesive (Ted Pella) was affixed to two electrode contact pads on the well slide. One pad was connected to ground, while the other to a function generator and high voltage amplifier, providing an electric potential of 200–600 V at 20–30 kHz to trap the sorted droplets in the wells by dielectrophoretic force.

**Printing procedure**

During the printing process, the printer head was fixed to an XYZ micromanipulator and the well substrate held on a motorized XY mechanical stage (MA-2000, ASI). The printing process is automated by custom LabView software which coordinates the droplet sorting of the printer head and the movement of the mechanical stage where the well substrate is held. When printing, the nozzle of the printer head was positioned close to the well substrate by the XYZ micromanipulator and the well substrate was moved across the wells in coordination with the sorting. After printing was complete, the bath oil was removed and the substrate dried by passive air convection. The substrate was then placed in a petri dish containing drying powders (Calcium chloride powder, Thermo Scientific, USA) and sealed with parafilm and stored at − 20 °C until needed.

**Matrix deposition and MALDI-Mass Spec imaging**

A matrix solution of 1,5-Diaminonaphthalene (DAN) (Millipore Sigma, USA) was prepared by adding 10 mg/mL DAN into a solution of 90% acetonitrile and 0.1% trifluoroacetic acid in DI water for the MS test of Naringenin and *Saccharomyces cerevisiae* yeast. It was loaded into an automated matrix sprayer (TM Sprayer, HTX Imaging) that coats the printed well substrate with a layer of DAN matrix at 30 °C, a flow rate of 0.125 mL/min, and velocity of 1200 mm/min. ~2 mL of matrix solution was used for each substrate. For MALDI MS imaging, the rapifleX TOF/TOF (Bruker, Germany) was used in negative ion mode at 50% laser power (4µJ), 200 shots and 50 µm pitch for the *Saccharomyces cerevisiae* yeast MS imaging. In a similar way, a matrix solution of Super-DHB (Millipore Sigma, USA) was used by adding 15 mg/mL into a solution of 90% acetonitrile and 0.1% trifluoroacetic acid in DI water for the MS test of TAL, naringenin, peptides, *Yarrowia lipolytica* yeast and mammalian cells. It is loaded into an automated matrix sprayer (TM Sprayer, HTX Imaging) that coats the substrate with a layer of

DHB matrix at 60 °C using a flow rate of 0.125 mL/min and velocity of 1200 mm/min. ~2 mL of matrix solution is used for each substrate. Solarix 7T (Bruker, Germany) is used in positive mode at 35% laser power, 300 shots and 80 μm pitch for TAL, naringenin and peptides test. rapifleX TOF/TOF (Bruker, Germany) is used in positive ion mode at 50% laser power (4 μJ), 200 shots and 50 μm pitch for the *Yarrowia lipolytica* yeast MS imaging.

**Mass Spec ion imaging analysis**

A Bruker flexImaging, SciLs Lab software (Bruker, Germany), and Cardinal 2 package in R[11] were used to analyze the mass spec imaging of the well substrate. The locations of target wells were identified and analyzed by a custom MATLAB program.

**Supplementary figures**

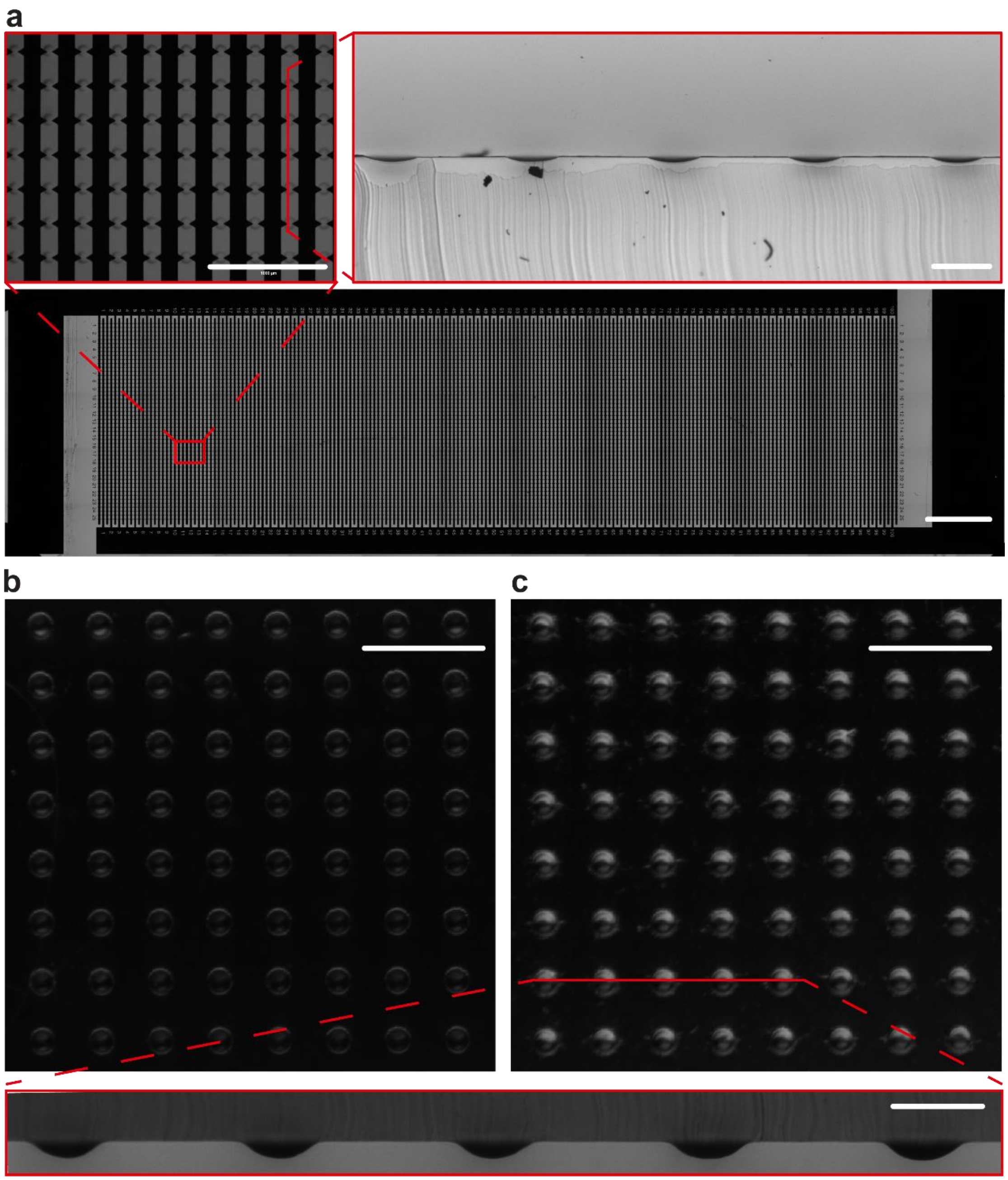


**Supplementary figure 1** | Different substrates with microbowls of various depth. **a,** Standard glass slide (75 by 25 mm) with 10,000 microbowls of a depth equals to 15.1µm ±2.3 µm. Scale bar is 5 mm. Top left insert is the enlarged view of the microbowls. Scale bar is 1 mm. Top right insert is the cross section view of the microbowls. Scale bar is 100 µm. **b**, and **c**, are the dark field image of microbowls. Scale bar is 300 µm. Bottom insert shows the cross section view of the microbowls in **c**. Scale bar is 100 µm.

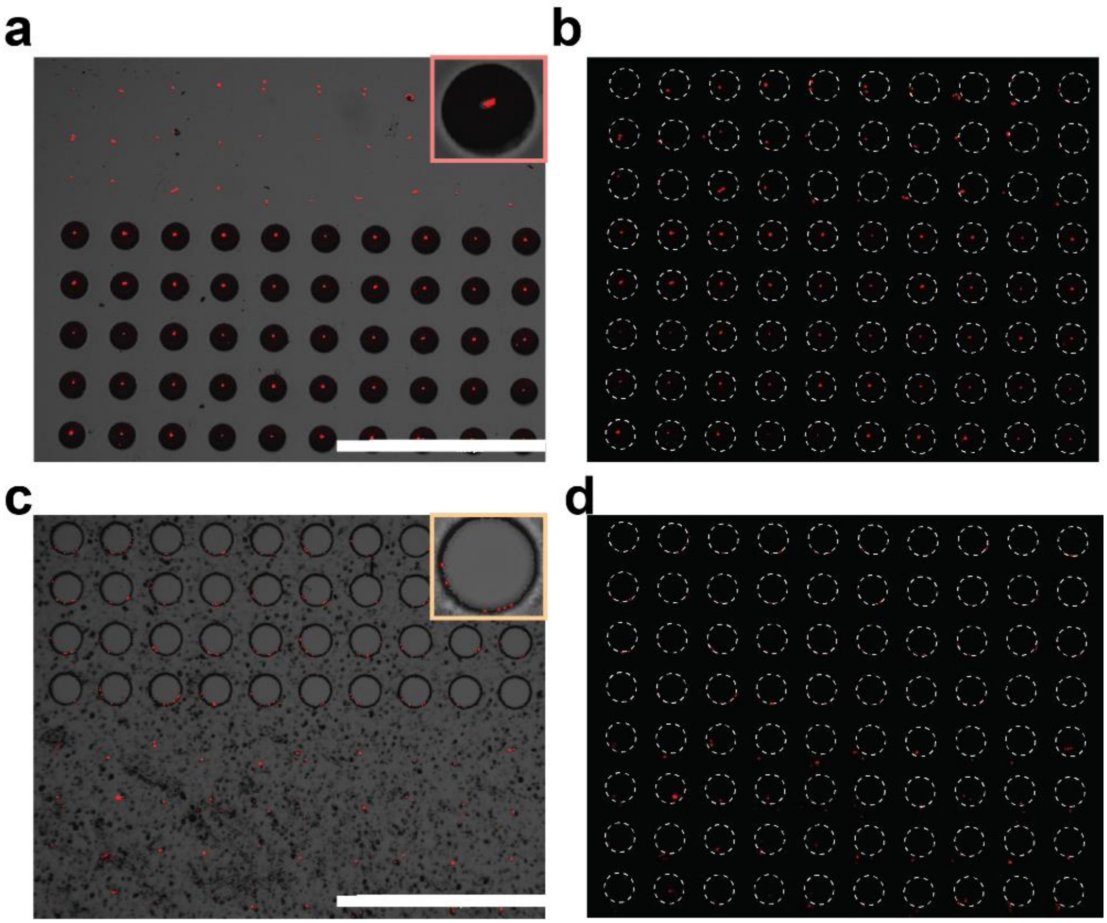


**Supplementary figure 2** | Different substrate with dried droplets containing beads for drying pattern analysis in Figure 3b. **a,** Droplets containing microbeads (red, RFP channel) were printed on flat and microbowls respectively, and then dried at room temperature. The top three rows are droplets printed on the flat surface and the bottom five row are droplets printed on microbowls. **b,** RFP epifluorescent image of **a**. Similar, **c,** shows the droplets printed on cylinder wells and flat surface respectively, and then dried at room temperature. **d,** RFP epifluorescent image of **c**. In **b** and **d,** the dotted white circles stand for the positions of the droplets where they are printed initially. Scale bars are 1 mm.

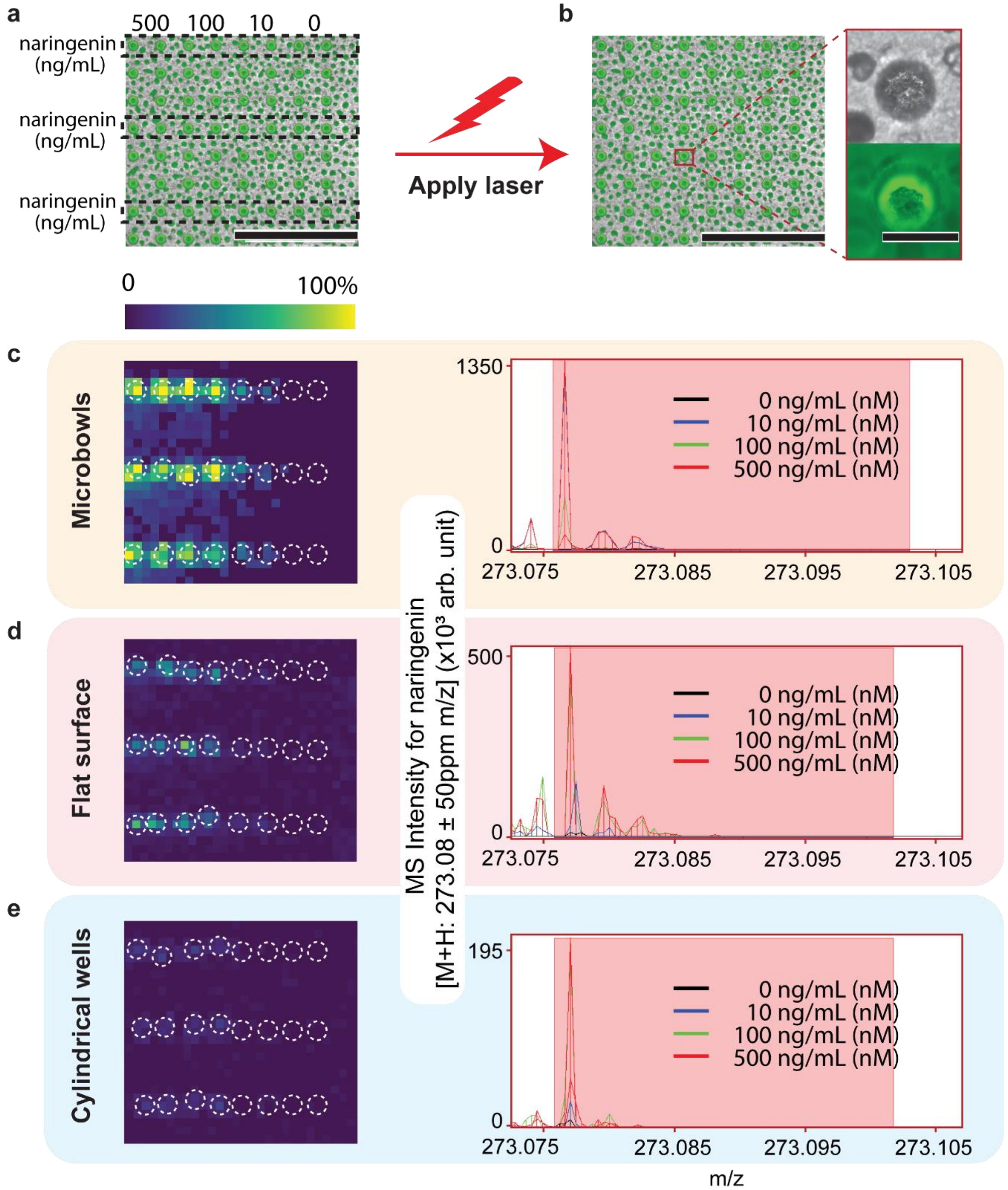


**Supplementary figure 3** | Different substrates with naringenin droplets for MS analysis in **Figure 4**. **a** and **b**, Combined images of transillumination and GFP epifluorescence channels of the microbowls, that were printed with droplets containing 500, 100, 10, and 0 ng/mL naringenin in water, dried and spray-coated with matrix, before and after laser scanning. Black dashed lines indicate where the droplets are printed. Insert image is the enlarged view of one microbowl after laser scanning. **c**, MALDI MS image of **a**, the microbowls. Right image is the enlarged spectrum for naringenin at different concentrations. Similar, **d** and **e**, shows flat surface and cylindrical wells respectively. White circles show which pixels are used for data analysis. Scale bars are 1 mm and 300 µm for the main figures and insert respectively.

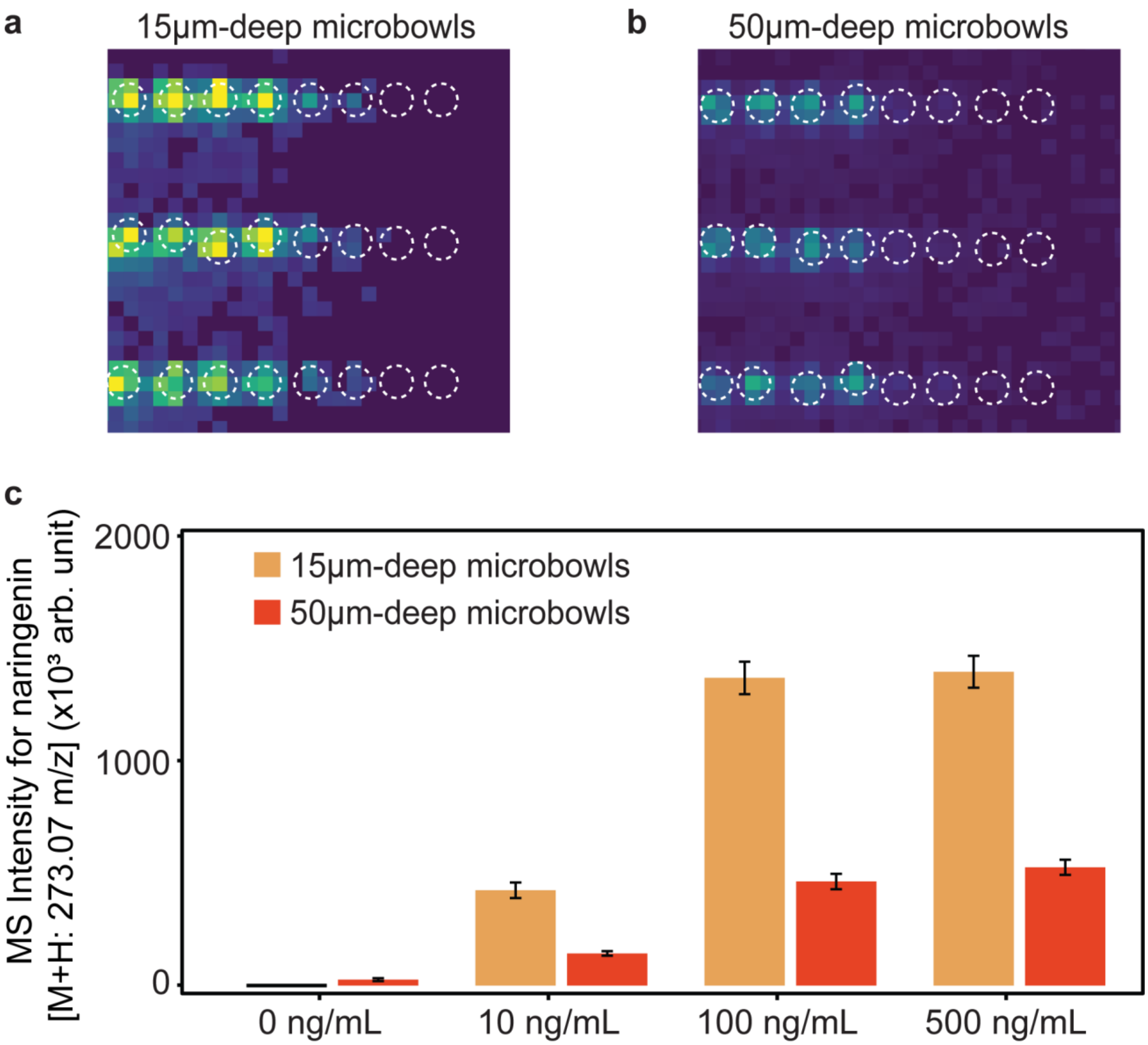


**Supplementary figure 4** | Microbowls with different depth for MS analysis of naringenin. **a** and **b**, MALDI MS image of microbowls with 15-µm and 50-µm depth respectively. Right image is the enlarged spectrum for naringenin at different concentrations. **c**, Sum of naringenin in MS signal for the microbowls with 15-µm and 50-µm depth respectively

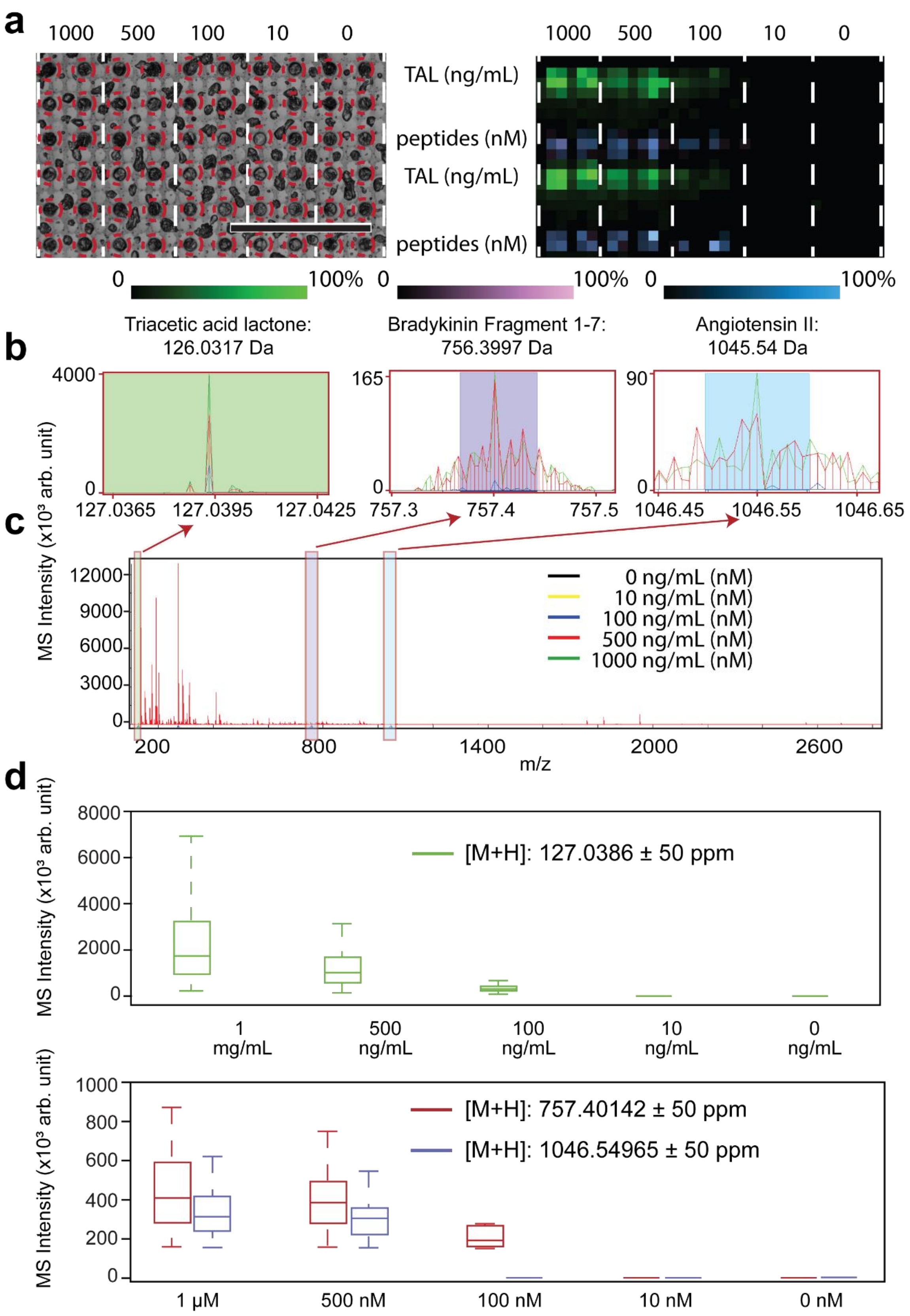


**Supplementary figure 5** | Microbowls with different analytes for MS analysis. **a**, Microbowls printed with different analytes after laser scanning, left, and the MS image of it, right. Red circles indicate where the microbowls are. **b**, enlarged spectrum for different analytes: green for TAL, purple and blue for peptides. **c**, the whole spectrum from all the microbowls. **d** and **e**, Sum of TAL, Bradykinin Fragment 1-7 and Angiotensin II peak amplitude in MS signal respectively. Scale bar is 1 mm.

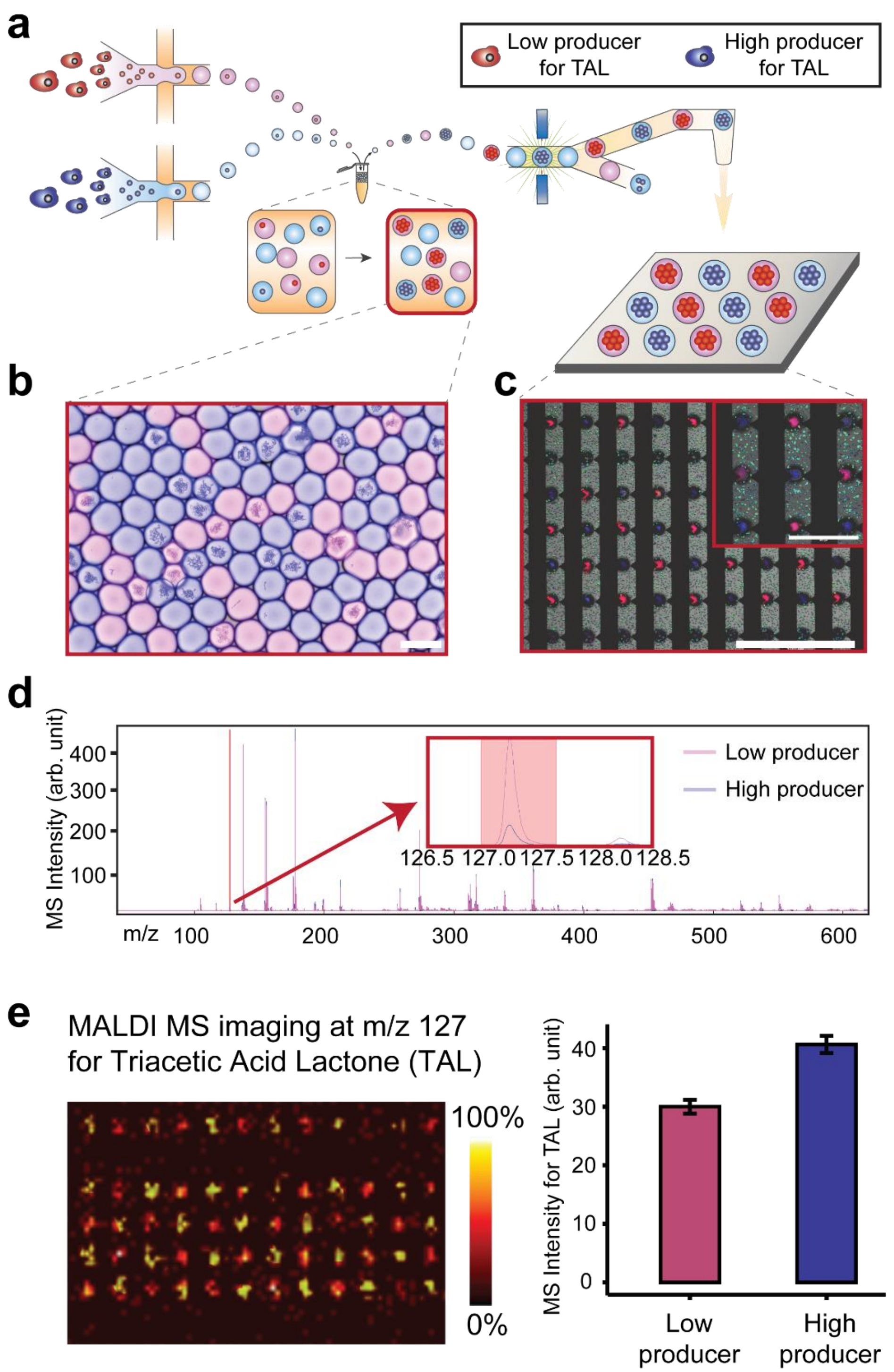


**Supplementary figure 6** | In-droplet coculture test of two strains of yeast that produce TAL. **a**, The two strains were encapsulated in droplet separately using the same protocol, then droplets were mixed together and printed to form a checkboard pattern. Red stands for low producer which produce around 0.3 mg/mL TAL and blue stands for high producer which produce around 0.8 mg/mL TAL. **b** shows how the droplets looks like after in-droplet coculture for 5 days. Scale bar is 100 µm. **c**, Combined image of Transillumination and RFP, DAPI epifluorescence channels of the dried droplets after applying matrix. Scale bars are 1mm and 300 µm for the main figure and insert respectively. **d** is the mass spectrum for the two strains of yeast with insert showing the enlarged spectrum for TAL. **e**, Left is the MALDI MS imaging for TAL and right is the sum of TAL in MS signal for the microbowls with low and high producer of TAL respectively.

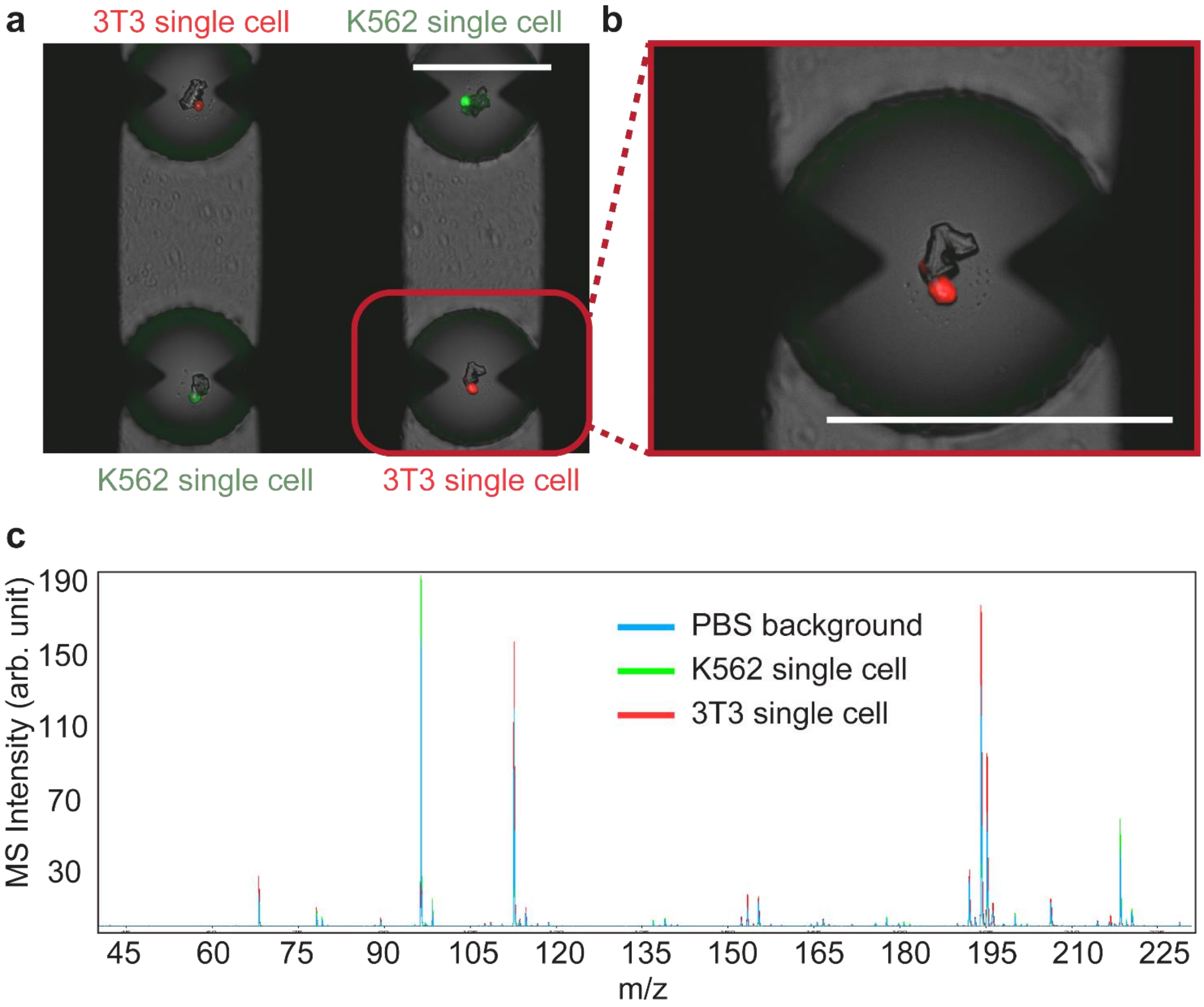


**Supplementary figure 7** | Microbowls with printed single cell for MS analysis. **a** Combined image of transillumination, GFP and RFP epifluorescence channels of the microbowls that are printed with single 3T3 and K562 cell per well. **b**, Enlarged image shows the crystal of salt and single 3T3 cell stained with red fluorescence. **c**, MS for the microbowls loaded with single K562, 3T3 cells and PBS droplet only, respectively.

**Video S1** Drying process of a water droplet containing microbeads on a flat surface.

**Video S2** Drying process of a water droplet containing microbeads on a cylinder well.

**Video S3** Drying process of a water droplet containing microbeads on a microbowl.

Reference for method section:

1. Nijkamp, J. F. *et al.* De novo sequencing, assembly and analysis of the genome of the laboratory strain Saccharomyces cerevisiae CEN.PK113-7D, a model for modern industrial biotechnology. *Microb Cell Fact* **11**, 36 (2012).
2. Chandran, S. & Shapland, E. Efficient Assembly of DNA Using Yeast Homologous Recombination (YHR). in *Synthetic DNA* (ed. Hughes, R. A.) vol. 1472 187–192 (Springer New York, 2017).
3. Ip, K., Yadin, R. & George, K. W. High-Throughput DNA Assembly Using Yeast Homologous Recombination. in *DNA Cloning and Assembly: Methods and Protocols* (eds. Chandran, S. & George, K. W.) 79–89 (Springer US, 2020). doi:10.1007/978-1-0716-0908-8_5.
4. Guthrie, C. & Fink, G. R. *Guide to Yeast Genetics and Molecular Biology*. (Gulf Professional Publishing, 2004).
5. Radford, A. Methods in yeast genetics — A laboratory course manual by M Rose, F Winston and P Hieter. pp 198. Cold Spring Harbor Laboratory Press, Cold Spring Harbor, New York. 1990. $34 ISBN 0-87969-354-1. *Biochemical Education* **19**, 101–102 (1991).
6. Griggs, D. W. & Johnston, M. Promoter elements determining weak expression of the GAL4 regulatory gene of Saccharomyces cerevisiae. *Molecular and Cellular Biology* **13**, 4999–5009 (1993).
7. Griggs, D. W. & Johnston, M. Regulated expression of the GAL4 activator gene in yeast provides a sensitive genetic switch for glucose repression. *PNAS* **88**, 8597–8601 (1991).
8. Markham, K. A. *et al.* Rewiring yarrowia lipolytica toward triacetic acid lactone for materials generation. *Proceedings of the National Academy of Sciences of the United States of America* **115**, 2096–2101 (2018).
9. Xia, Y. & Whitesides, G. M. SOFT LITHOGRAPHY. *Annual Review of Materials Science* **28**, 153–184 (1998).
10. Cole, R. H. *et al.* Printed droplet microfluidics for on demand dispensing of picoliter droplets and cells. *Proceedings of the National Academy of Sciences of the United States of America* **114**, 8728–8733 (2017).
11. Bemis, K. D. *et al.* Cardinal: An R package for statistical analysis of mass spectrometry-based imaging experiments. *Bioinformatics* **31**, 2418–2420 (2015).
